# PhysiPKPD: A pharmacokinetics and pharmacodynamics module for PhysiCell

**DOI:** 10.1101/2022.09.12.507681

**Authors:** Daniel Bergman, Lauren Marazzi, Mukti Chowkwale, Deepa Maheshvare M, Supriya Bidanta, Tarunendu Mapder, Jialun Li

## Abstract

Pharmacokinetics and pharmacodynamics are key considerations in any study of molecular therapies. It is thus imperative to factor their effects in to any *in silico* model of biological tissue involving such therapies. Furthermore, creation of a standardized and flexible framework will benefit the community by increasing access to such modules and enhancing their communicability. PhysiCell is an open source physics-based cell simulator, i.e. a platform for modeling biological tissue, that is quickly being adopted and utilized by the mathematical biology community. We present here PhysiPKPD, an open source PhysiCell-based package that allows users to include PKPD in PhysiCell models.

**Availability & Implementation:** The source code for PhysiPKPD is located here: https://github.com/drbergman/PhysiPKPD.

## Statement of Need

Agent-based modeling has become a common tool for research in systems biology. However, it is still a relatively new tool and the community remains in need of standardized models that promote accessibility and extensibility. The PhysiCell platform is one such effort to achieve this [1, 2]. It is open-source, actively maintained, and has taken large steps towards being widely adopted as evidenced by the release of PhysiBoSS, an independently-developed integration of PhysiCell with intracellular signaling using Boolean modeling. New add-ons to PhysiCell will further broaden its appeal and facilitate the use of standards that will achieve the aforementioned community goals of accessibility and extensibility.

Many agent-based models (ABMs) have been developed to study diseases, including cancer [3, 4, 5, 6, 7, 8], COVID-19 [9], tuberculosis [10, 11], and more, with a core purpose being to determine the best means to treat a patient with the given disease. This involves modeling therapeutics alongside the cells in a given microenvironment. There are two key pieces of including drugs: how much gets to the cells and what they then do to the cells. These two pieces are referred to as pharmacokinetics and pharmacodynamics, respectively. Together, they are often called PKPD.

The dynamics involved in PKPD are generalizable to many different drugs acting in many different ways. This makes it an appealing target for a single module that can flexibly handle these processes on a platform such as PhysiCell. Features on both the PK side–such as dosing schedules, loading doses, elimination and distribution rates–and the PD side–such as mechanisms of action (MOA), effect, EC50, and hill coefficients–are amenable to this level of abstraction.

We present here PhysiPKPD, a standardized framework to incorporate these PKPD processes in PhysiCell. We also provide several examples demonstrating PhysiPKPD and providing users with two template projects to aid in the model-building process. In this first version, we provide the three options for modeling PK dynamics: 1-compartment and 2-compartment models with linear clearance rates and the ability to supply an SBML-defined model. On the PD side, substrates cause damage to cells based on the internalized substrate. As this damage accumulates within a cell, the associated MOA-associated rate parameter(s) tend towards a user-defined saturation rate. Both PK and PD dynamics can be added independently to any substrate in the simulation. Substrates can target any cell type with PD dynamics.

## Implementation

We provide two ways to create and run PhysiPKPD models, shown in Figure 2. First, the sample projects that come with PhysiPKPD demonstrate the MOAs implemented in PhysiPKPD (Figure 2A). Second, the user can use the template projects to jumpstart the model-building process with the full range of possible parameters and code present throughout the PhysiCell repository (Figure 2B). We now explain how PhysiPKPD achieves these dynamics. In what follows, `S` will stand for the name of a substrate, `C` will stand for the name of a cell type, and X will stand for a MOA.

**Figure 1.**
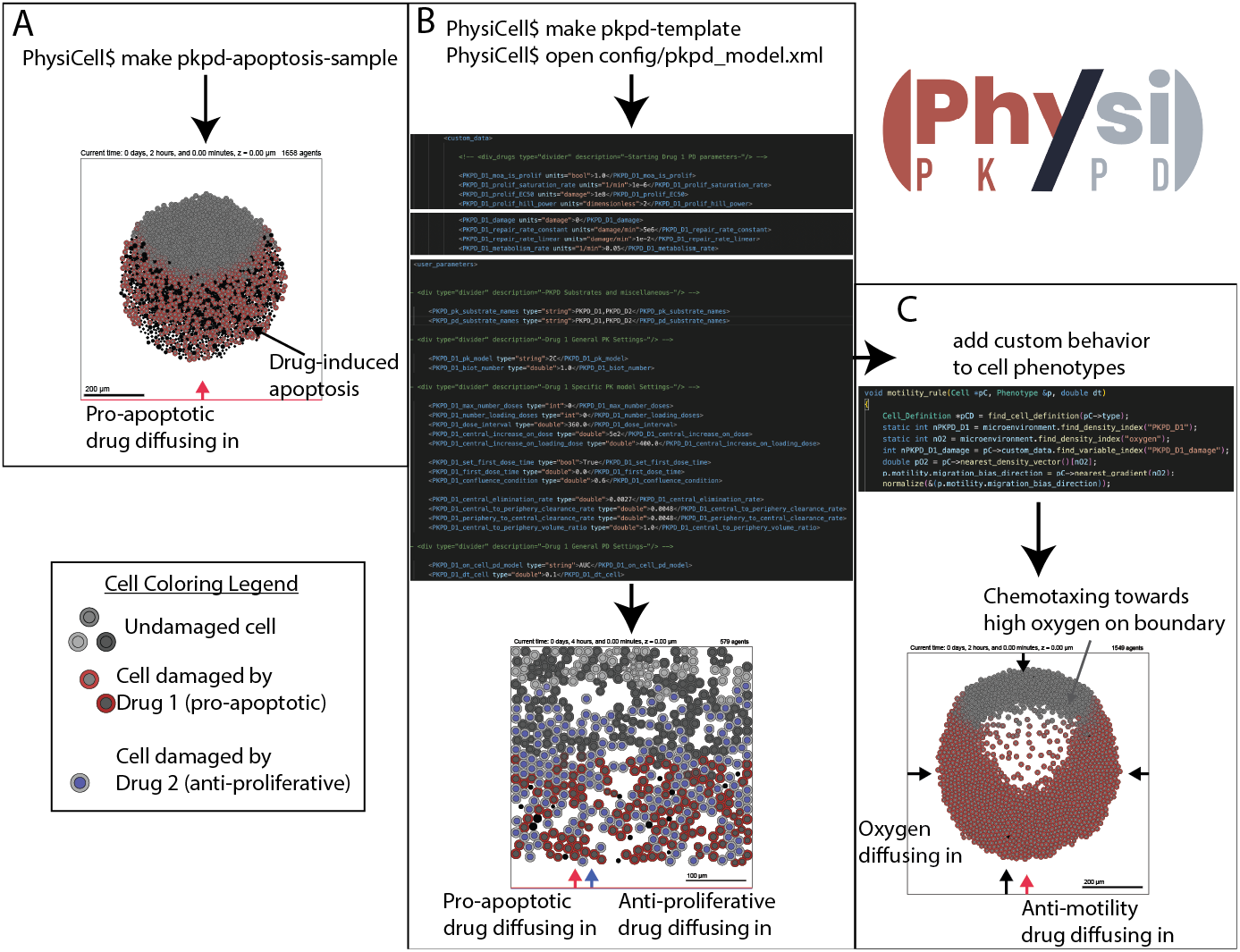
How to implement PhysiPKPD. A: Using the sample projects that come with PhysiPKPD. B: Using the model builder to PhysiPKPD modifications to the model builder to create a wide variety of models. C: Editing the C++ code in custom modules to further refine the model, e.g. making cell chemotax along an oxygen gradient.

**Figure 2.**
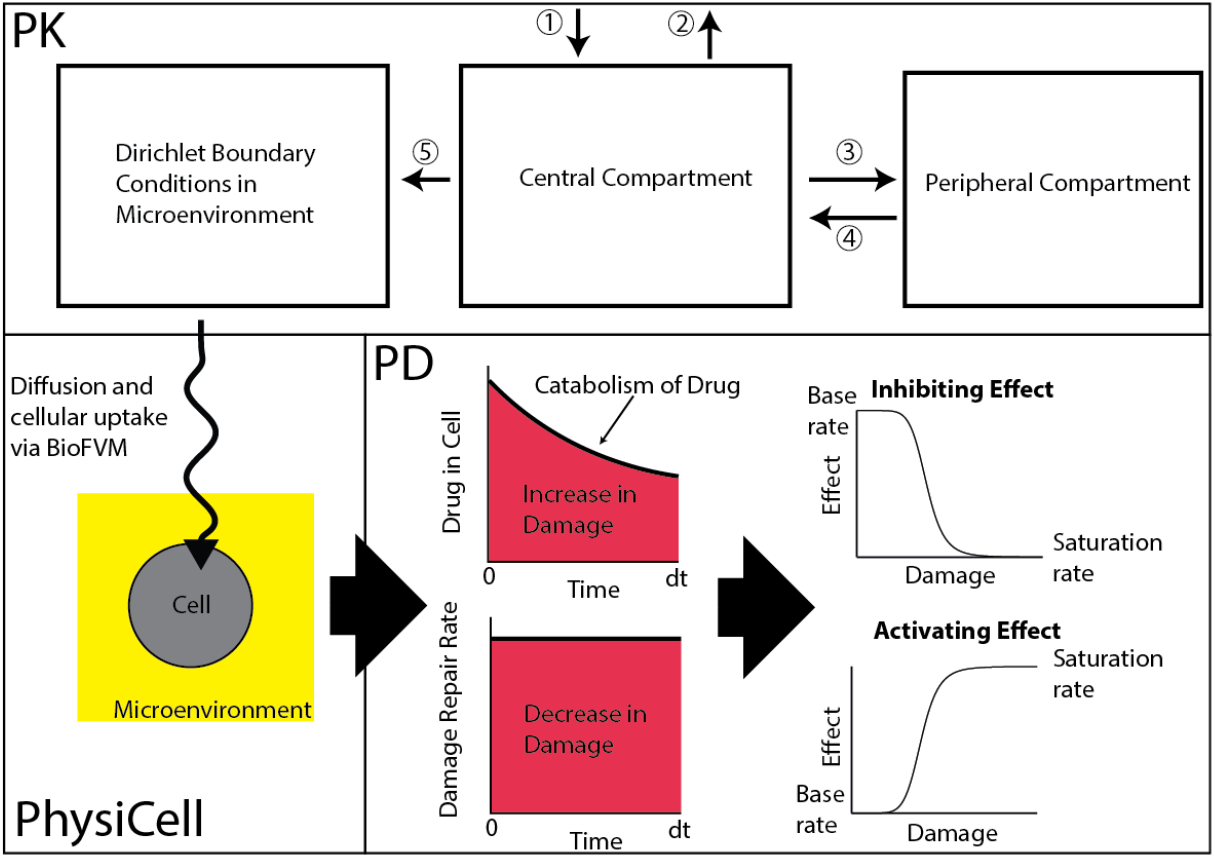
Cartoon of integration of PK and PD modules with PhysiCell. PK: (1) Administration of drug in central compartment. (2) Elimination of drug. (3) Distribution of drug into peripheral compartment. (4) Redistribution of drug back into central compartment. (5) Extravasation of drug into microenvironment. PhysiCell: Diffusion and cellular uptake of drug in the microenvironment. PD: Drug causes damage to cells over time. Cells can catabolize the drug and repair the damage. Damage determines the effect of the drug on its MOA.

## Pharmacokinetics

For pharmacokinetics, the user must first specify which substrates in the model follow PK dynamics. This is done by adding these substrate names to the comma-separated string user parameter `PKPD_pk_substrate_names`.

### PK models

For each of these substrates, the user should specify a PK model (see Table 1). If this is not specified, PhysiPKPD will attempt to use a two-compartment model. In addition, the user should also specify a Biot number, which PhysiPKPD uses to set the ratio of perivascular substrate concentration to that in the blood vessel [12]. PhysiPKPD will default to a value of 1, indicating equal concentrations inside and outside of the capillary walls.

**Table 1.** PK model specifications. The specification column indicates the string value to be used in the configuration file for the parameter `S_pk_model`.

| Model | Description | Specification |
| --- | --- | --- |
| One-compartment | Circulation compartment with linear elimination | 1C |
| Two-compartment | Circulation and peripheral compartments with linear clearance rates | 2C |
| SBML-defined | Any SBML-defined model with one species named `circulation_concentration` | SBML |

For example, add the following to User Parameters in the configuration file: `<S_pk_model type="string">2C</S_pk_model>`

### One-compartment models

The PhysiPKPD one-compartment model obeys the differential equation

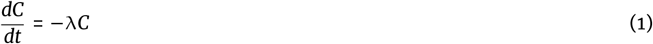

where *C* represents the circulation concentration of the substrate and λ is the elimination rate (see Table 2). PhysiPKPD uses the value of *C* to update the Dirichlet conditions in the PhysiCell microenvironment after multiplying by the Biot number.

**Table 2.** PK parameters for 1- and 2-compartment models.

| Parameter | Description | Type | Units |
| --- | --- | --- | --- |
| `S_central_elimination_rate` ($\lambda$) | Rate of elimination of the drug in systemic circulation | double | $\text{min}^{-1}$ |
| `S_central_to_periphery_volume_ratio` * ($R$) | Ratio of volumes of central and peripheral compartments to conserve mass of drug during distribution and redistribution | double | None |
| `S_central_to_periphery_clearance_rate` * ($k_{12}$) | Rate of change in concentration in central compartment due to distribution | double | $\text{min}^{-1}$ |
| `S_periphery_to_central_clearance_rate` * ($k_{21}$) | Rate of change in concentration in periphery compartment due to redistribution | double | $\text{min}^{-1}$ |

Replace *s* with the name of the substrate.
\*Parameter is not necessary for 1-compartment models.

### Two-compartment models

The PhysiPKPD two-compartment model obeys the differential equation

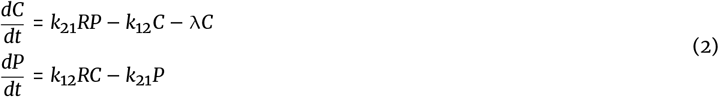

Here, *C* and λ are as above. *P* is the periphery concentration. The parameters *k*_12_ and *k*_21_ are the intercompartmental clearance rates and *R* is the ratio of the volumes of the central and peripheral compartments (see Table 2). PhysiPKPD uses the value of *C* to update the Dirichlet conditions in the PhysiCell microenvironment after multiplying by the Biot number. The inclusion of the periphery compartment allows for biphasic elimination in the central compartment.

### SBML-defined models

If the above two models are inadequate for a user’s purposes, an SBML file can be used to specify a PK model. We have used Copasi [13] to build such models, but any program that outputs an SBML file will work, e.g., sbmlutils [14]. PhysiPKPD will use the first state variable of the model to update Dirichlet conditions in the PhysiCell microenvironment after multiplying by the Biot number.

### Dosing schedules

For the one- and two-compartment models, all parameters and dosing events must be specified within the configuration file (see Table 3). For any missing parameters, PhysiPKPD will issue warnings and use default values where it can, and it will throw errors where it must. All substrates are assumed to be given intravenously so that the concentration in the central compartment has a one-time increase upon dosing, `S_central_increase_on_dose`. These doses are given on regular intervals, `S_dose_interval`, until either the simulation ends or until the maximum number of doses has been administered. The first dose can be given at a fixed time or based on confluence in 2D in the entire rectangular microenvironment. A loading dose can also be set with a fixed number of doses. Future releases can include other methods of administration, e.g. oral, and also include finer-grained control such as more sophisticated timings for doses, e.g. M-F dosing.

**Table 3.**
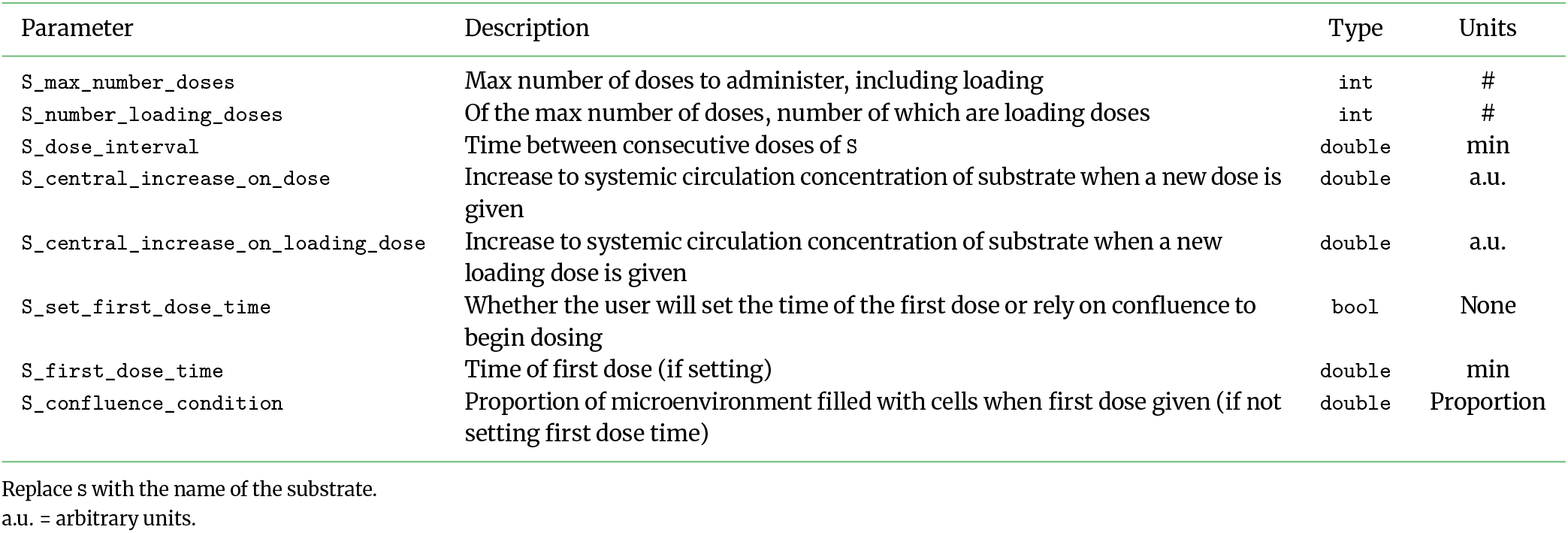
Dosing parameters for 1- and 2-compartment models.

For SBML-defined models, the user must include dosing events in the SBML file itself. In Copasi, this can be done by creating Events that increase the concentration of a compartment(s) at certain times. In the future, we hope to allow the user to specify a CSV file with the dosing times and amounts for a substrate along with an SBML-defined system of ODEs (without dosing events) that PhysiPKPD then combines into a new SBML with dosing events.

## Pharmacodynamics

Analogous to identifying substrates following PK dynamics, the user must specify which substrates follow PD dynamics. This is done by adding these substrate names to the comma-separated string user parameter `PKPD_pd_substrate_names`. The list of PK substrates and the list of PD substrates need not have any relationship.

### Damage accumulation

For each cell type affected by a particular substrate, a damage-accumulation model should be specified. If this is not specified, PhysiPKPD will default to the concentration-based model. The two options are the concentration-based model and the amount-based model (see Table 4). Both of these models use the area under the curve (AUC) of the internalized substrate to determine the current damage by the substrate on the cell. Future releases of PhysiPKPD may allow for SBML-defined damage accumulation functions. The experienced PhysiCell user can opt to use the already-implemented Intracellular features of PhysiCell to achieve this already.

**Table 4.** Damage accumulation model specifications. The specification column indicates the string value to be used in the configuration file for the parameter S_on_C_pd_model.

| Model | Description | Specification |
| --- | --- | --- |
| Concentration-based | Damage accumulates in a cell based on the concentration of the internalized substrate | AUC |
| Amount-based | Damage accumulates in a cell based on the amount of the internalized substrate | AUC_amount |

For example, add the following to User Parameters in the configuration file: `<S_on_C_pd_model type="string">AUC</S_on_C_pd_model>`

For either model, let *A* represent the internalized substrate, concentration or amount. Cells metabolize the substrate at a rate *m*. Cells then accumulate damage proportional to the internalized substrate. This damage is repaired at a linear rate *r*_1_ and a constant rate of *r*_0_. As damage is an abstract quantity, the proportionality constant for the internalized substrate causing damage is set to 1. All three of these parameters (Table 5) must be included in the custom data for `C`, the cell type affected by substrate `S`. The differential equations are then given by

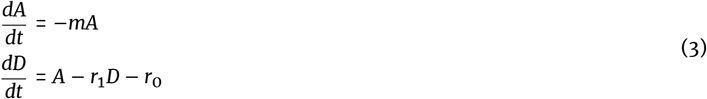

**Table 5.** Damage accumulation parameters.

| Parameter | Description | Units |
| --- | --- | --- |
| $S_{\text{metabolism\_rate}} (m)$ | Metabolism rate of internalized substrate | $\text{min}^{-1}$ |
| $S_{\text{repair\_rate\_constant}} (r_0)$ | Zero-order repair rate of damage | damage/min |
| $S_{\text{repair\_rate\_linear}} (r_1)$ | First-order repair rate of damage | $\text{min}^{-1}$ |

Set all in Custom Data of the cell definition of c.

This equation is solved by default at the mechanics time step. The user can change this by setting the parameter S_dt_C. PhysiPKPD uses the analytic solution to solve these dynamics. By default, PhysiPKPD does this by pre-computing the relevant quantities. However, in cases where the above parameters can vary by cell type, the user should set the parameter `PKPD_precompute_all_pd_quantities` to false.

Finer-grained control can be set by S_precompute_pd_for_C, which overrides `PKPD_precompute_all_pd_quantities` on a case-by-case basis.

### Cell effect parameters

To apply the desired MOA of `S` on `C`, the damage variable `S_damage` is used as the input to a Hill-type function. The user must specify three parameters for this Hill-type function in addition to identifying this MOA (see Table 6). The four MOAs currently implemented in PhysiPKPD are proliferation (prolif), apoptosis (apop), necrosis (necrosis), and motility (motility). Replace X in Table 6 with the parenthetical name of the desired MOA. By default, all MOAs are assumed off, so including `S_moa_is_X` in the custom data is only necessary if set to true. Note that custom data in PhysiCell must be of type double, so “true” means a value > 0.5.

**Table 6.** Cell effect parameters.

| Parameter | Description | Units |
| --- | --- | --- |
| $S_{\text{moa\_is\_X}}$ | Identifies whether the MOA of s on c is X | None (Values > 0.5 trigger the MOA) |
| $S_{\text{X\_saturation\_rate}}$ | Limiting rate of X as damage from s grows towards infinity | $\text{min}^{-1}$ |
| $S_{\text{X\_EC50}}$ | Damage from s at which the rate of X is halfway between the base and saturation rates | damage |
| $S_{\text{X\_hill\_power}}$ | Hill power | None |

Replace X with one of the following: prolifer, apopt, necrosis, or motility.

### Multiply-targeted MOAs

In the case that two or more substrates have the same MOA on a given cell type, we compute the drug effects as multipliers or factors. That is, the algorithm computes the saturation factor, *f*_sat_, based on the user-supplied saturation rate and uses the damage to compute a factor, *f*, between 1 (no change) and the saturation factor (max change):

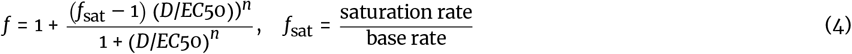

Then, when multiple substrates affect proliferation, for example, the factors for each are multiplied to give the final factor for the cell. In the case of necrosis, it is often the case that the base rate for necrosis is set to 0, which causes problems in computing *f*_sat_ above. Thus, we instead add the effects of multiple substrates targeting necrosis.

### Resetting to base rates

The implementation of PD dynamics is often done in the context of other dynamics. Typical examples used in the PhysiCell community include extracellular oxygen concentration and overcrowding (sometimes determined by simple pressure). PhysiPKPD therefore does not reset the MOA-targeted rates to their base value before applying the output of the Hill-type function. Thus, it is incumbent on the user to make sure the effects on these rates do not stack each time these effects are computed. See the `custom.cpp` files in any of the PhysiPKPD sample projects or the template projects to see how this can be done.

## Example Results

We provide several examples of implementations of PhysiPKPD: one for each MOA, one combination therapy with one anti-proliferative drug and one pro-apoptotic drug (Figure 1A), and one using a confluence condition to start dosing. Follow the README.md file instructions to get PhysiPKPD set up in the root PhysiCell directory. The output will go to the output folder. There are many parameters that can be changed to explore various behaviors, even restricted to just one MOA. See Tables 2, 3, 5, and 6 for parameters and their descriptions that are unique to PhysiPKPD. In Figure 3, we show results from the anti-proliferative drug sample project, including creating an IC50 curve by varying the dose over several orders of magnitude.

**Figure 3.**
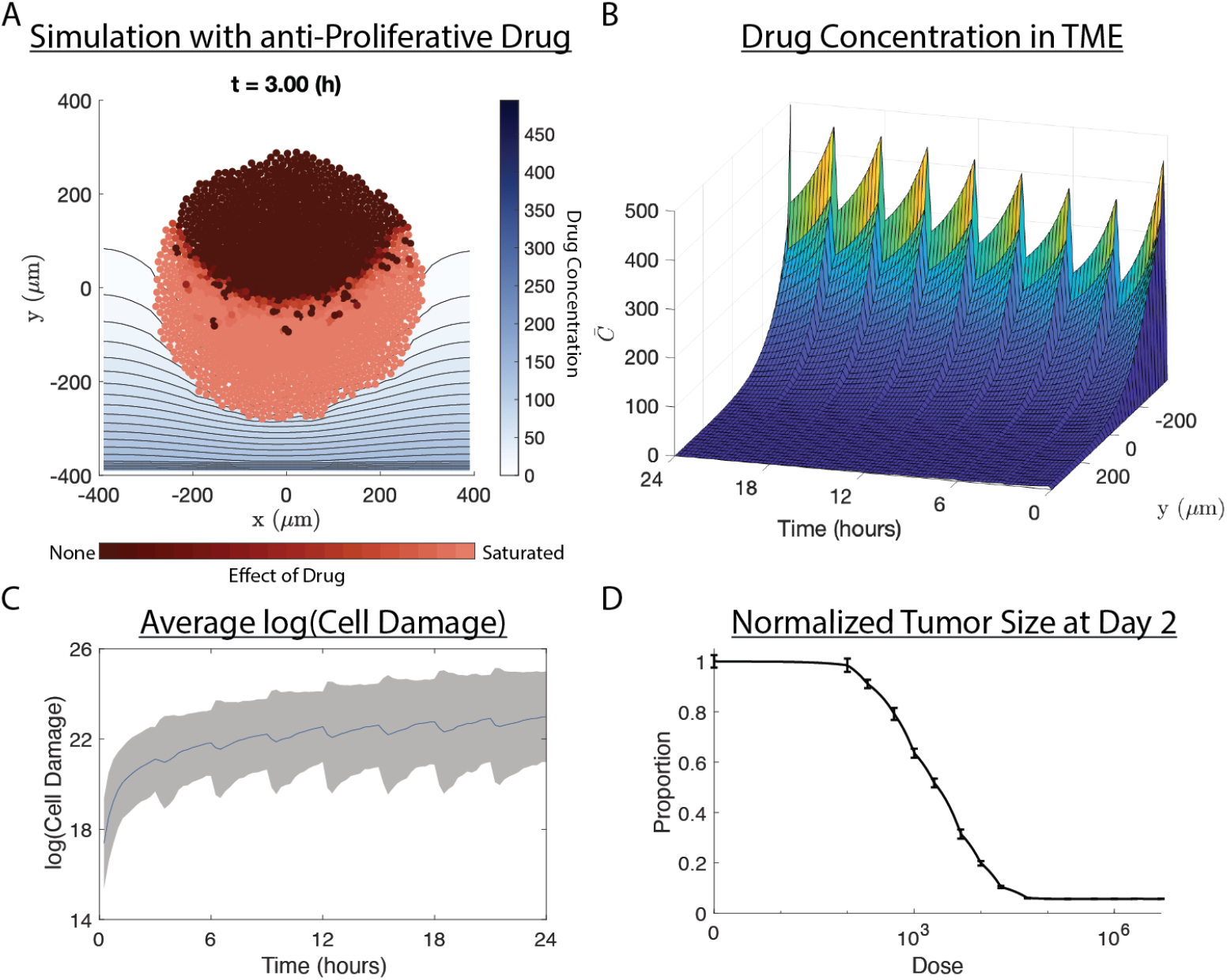
Results of PhysiPKPD moa_proliferation example. An anti-proliferative drug enters the microenvironment from the bottom boundary (y=-400 µm) and diffuses up towards the tumor. A: Snapshot of the simulation after 3 hours. Drug concentration shown by contour plot. Cells colored by the current effect of drug damage on their proliferation rates. B: Drug concentration in the microenvironment over time. The average is taken over the x dimension. C: Average logarithm of cell damage over time. Average is taken only over cells with positive damage. Gray area represents +/- SD of the log-transformed damage. D: IC50 curve generated from running this example with just one dose of drug given at time t=0 over 16 different doses including a control with no drug. Mean and SD for each dose computed from 5 samples. Tumor size at t=48 hours relative to the control shown. Note: analysis and initial plotting of A-D were done in MATLAB using the output from the PhysiCell simulation. The code that that produces these figures is not part of PhysiPKPD.

## Discussion

A PKPD model that is ready to use “out of the box” with PhysiCell will greatly benefit those seeking to use PhysiCell as a platform for agent-based modeling. It will also create a standard which can be applied across any modelers looking to incorporate PKPD dynamics into their PhysiCell models.

There are many improvements and additions we hope to make to PhysiPKPD in the near future. With regards to PK, we hope to make it easier to include dosing events in an SBML file by allowing the user to supply a PK model and a CSV with the dosing information. With regards to PD, there are many features that can be added to allow for greater flexibility in terms of mechanisms of action, effect models, and integration with intracellular signaling. We have so far assumed that the AUC of a substrate inside a given cell, what we called damage, can be used to determine the effect of a drug on that cell. This is similar to irreversible effects models which use the AUC of the drug concentration to determine the response [15]. Many other effects models have been used, including simple direct effect, indirect response, and signal transduction models [16]. Recent work has used Bayesian inference to determine the distribution of delay times in response to therapeutic agents [17], making stochastic effects models an appealing next step as well. This could be added directly into previously-studied intracellular models or be used as a means to coarse-grain such complex models while still allowing for heterogeneity within cell types.

We are also working with the makers of PhysiCell to further simplify and standardize the integration of PhysiPKPD into PhysiCell. A key goal of this partnership will be to allow the user to include PKPD dynamics while only needing to specify the relevant parameters and not directly interact with the C++ code.

We look forward to continuing to develop this tool as the community uses it and seeks new features.

## Availability of source code and requirements

- Project name: PhysiPKPD
- Project home page: https://github.com/drbergman/PhysiPKPD
- Operating system(s): Platform independent
- Programming language: C++
- Other requirements: PhysiCell 1.10.4 or higher, libRoadRunner
- License: e.g. BSD 3-clause license

## Declarations

## List of abbreviations

ABM: agent-based model
MOA: mechanism-of-action
PD: pharmacodynamics
PK: pharamcokinetics

## Ethical Approval (optional)

Not applicable

## Author’s Contributions

Conceptualization: DB, LM, MC, TM; Methodology: DB; Software: DB, LM, MC, DMM, SB; Validation: DB, LM, MC, DMM, SB, TM, JL; Visualization: DB; Writing–original draft: DB; Writing–review & editing: DB, LM, MC, DMM, SB, TM, JL

## Acknowledgements

We would like to thank the PhysiCell team for all the help they have provided along the way. In particular, we would like to thank Paul Macklin, Randy Heiland, John Metzcar and Furkan Kurtoglu.

